# Do Muscle Synergies Improve Optimization Prediction of Muscle Activations During Gait?

**DOI:** 10.1101/851873

**Authors:** Florian Michaud, Mohammad S. Shourijeh, Benjamin J. Fregly, Javier Cuadrado

## Abstract

Determination of muscle forces during motion can help to understand motor control, assess pathological movement, diagnose neuromuscular disorders, or estimate joint loads. Difficulty of in vivo measurement made computational analysis become a common alternative in which, as several muscles serve each degree of freedom, the muscle redundancy problem must be solved. Unlike static optimization (SO), synergy optimization (SynO) couples muscle activations across all time frames, thereby altering estimated muscle co-contraction. This study explores whether the use of a muscle synergy structure within a static optimization framework improves prediction of muscle activations during walking. A motion/force/EMG gait analysis was performed on five healthy subjects. A musculoskeletal model of the right leg actuated by 43 Hill-type muscles was scaled to each subject and used to calculate joint moments, muscle-tendon kinematics and moment arms. Muscle activations were then estimated using SynO with two to six synergies and traditional SO, and these estimates were compared with EMG measurements. SynO neither improved SO prediction of experimental activation patterns nor provided SO exact matching of joint moments. Finally, synergy analysis was performed on SO estimated activations, being found that the reconstructed activations produced poor matching of experimental activations and joint moments. As conclusion, it can be said that, although SynO did not improve prediction of muscle activations during gait, its reduced dimensional control space could be beneficial for applications such as functional electrical stimulation (FES) or motion control and prediction.

## 1. Introduction

Knowledge of muscle forces during human movement could elucidate basic principles of human motor control (M.R. Pierrynowski and Morrison 1985), facilitate assessment of pathological movement and diagnosis of neuromuscular disorders, and improve estimation of the loads experienced by diseased or injured joints (Hardt 1978). Because in vivo measurement of muscle force is invasive and impossible for some muscles, computer modeling has become a commonly used alternative approach (Nagano et al. 2005). However, because more muscles than degrees of freedom exist in the human musculoskeletal system, an infinite number of recruitment patterns are possible mathematically. This problem is often referred to as the muscle redundancy problem (Damsgaard et al. 2006) or force-sharing problem (Dul et al. 1984).

The muscle redundancy problem is commonly solved by an inverse dynamic optimization method called Static Optimization (SO) (Ambrósio and Kecskeméthy 2007; Crowninshield 1978; Shourijeh, Mehrabi, and McPhee 2017), which considers muscle activations as if each muscle was activated independently. However, recent studies have demonstrated that the central nervous system (CNS) appears to use muscle synergies to simplify neural control of movement by coupling muscle activations together (Merkle et al. 1998)(Shourijeh, Flaxman, and Benoit 2016)(Barroso et al. 2017). Synergies take a high dimensional control space and reduce it to a low dimensional space, which is potentially useful for reducing the level of indeterminacy when estimating muscle forces via optimization. Several recent studies have used a synergy structure to reduce the dimensionality of the unknown muscle activation controls (McGowan et al. 2010; Neptune, Clark, and Kautz 2009; Mehrabi, Schwartz, and Steele 2019). However, the models used in these studies were limited to sagittal plane motion and used a reduced number of muscles because the synergy information was extracted from EMG measurements available from only superficial muscles.

Recently, a computational approach called Synergy Optimization (SynO) has been proposed that uses muscle synergies to reduce indeterminacy when estimating leg muscle forces during walking (S. Shourijeh and Fregly 2019). The authors evaluated how the specified number of synergies affected estimated lower body joint stiffness and inverse dynamic joint moment matching. While results obtained from SynO were compared with those obtained from SO, experimental evaluation of the muscle activations predicted by SynO was not performed. Furthermore, because imposition of a synergy structure on predicted muscle activations ties all time frames together, SynO is more complex and slower computationally than is SO. Unlike previous approaches that used sagittal plane models with a reduced number of muscles, SynO allows the use of more complex and realistic musculoskeletal models to estimate full leg muscle synergies and corresponding muscle activations.

This study evaluated whether imposition of a synergy structure on muscle activations estimated via inverse dynamic optimization (i.e., SynO) produces muscle activation estimates that are more consistent with EMG measurements than are those produced by traditional SO. Muscle activations reconstructed by performing synergy analysis on SO activations were included in the evaluation as well. Muscle activations and inverse dynamic joint moment matching from all three approaches were compared to activations derived from experimental EMG data and joint moments calculated by inverse dynamics using data collected from five subjects performing overground walking. Three-dimensional models of the subjects were used to perform the evaluation. Comparison of these three approaches provides insight into the extent to which, and the conditions under which, imposition of a synergy structure may improve the estimation of muscle forces during walking.

## 2. Methods

### 2.1 Experimental data collection

Five subjects (four males, one female, age 42 ± 16 years, height 178 ± 11 cm, body mass 75 ± 25 kg) were recruited for this study. All subjects gave written informed consent for their participation. Subjects walked at their self-selected speed (1.1 ± 0.18 m/s) along a walkway with two embedded force plates (AMTI, AccuGait sampling at 100 Hz). The motion was captured using 12 optical infrared cameras (Natural Point, OptiTrack FLEX:V100 also sampling at 100 Hz) that computed the position of 37 optical markers. Additionally, 11 surface EMG signals on the right leg were recorded at 1 kHz (BTS, FREEEMG). Each EMG signal was rectified, filtered by singular spectrum analysis (SSA) with a window length of 250 (Romero et al. 2015) (equivalent to the common forward and reverse low-pass 5th order Butterworth filter with a cut-off frequency of 15 Hz) and then normalized with respect to its maximal value as recommended in (Raison et al. 2011).

### 2.2 Musculoskeletal model creation

The human body was modeled as a three-dimensional multibody system formed by rigid bodies (Figure 1, left and center). The model consisted of 18 anatomical segments (Lugrís, Carlín, Pàmies-Vilà, et al. 2013): two hindfeet, two forefeet, two shanks, two thighs, a pelvis, a torso, a neck, a head, two arms, two forearms, and two hands. The segments were linked by ideal spherical joints, thus defining a model with 57 degrees of freedom (DOFs). The axes of the global reference frame were defined as follows: x-axis in the anterior–posterior direction, y-axis in the medial–lateral direction, and z-axis in the vertical direction. The computational model was defined with 228 mixed (natural + angular) coordinates. The subset of natural coordinates comprised the three Cartesian coordinates of 22 points and the three Cartesian components of 36 unit vectors, thus yielding a total of 174 variables.

**Figure 1:**
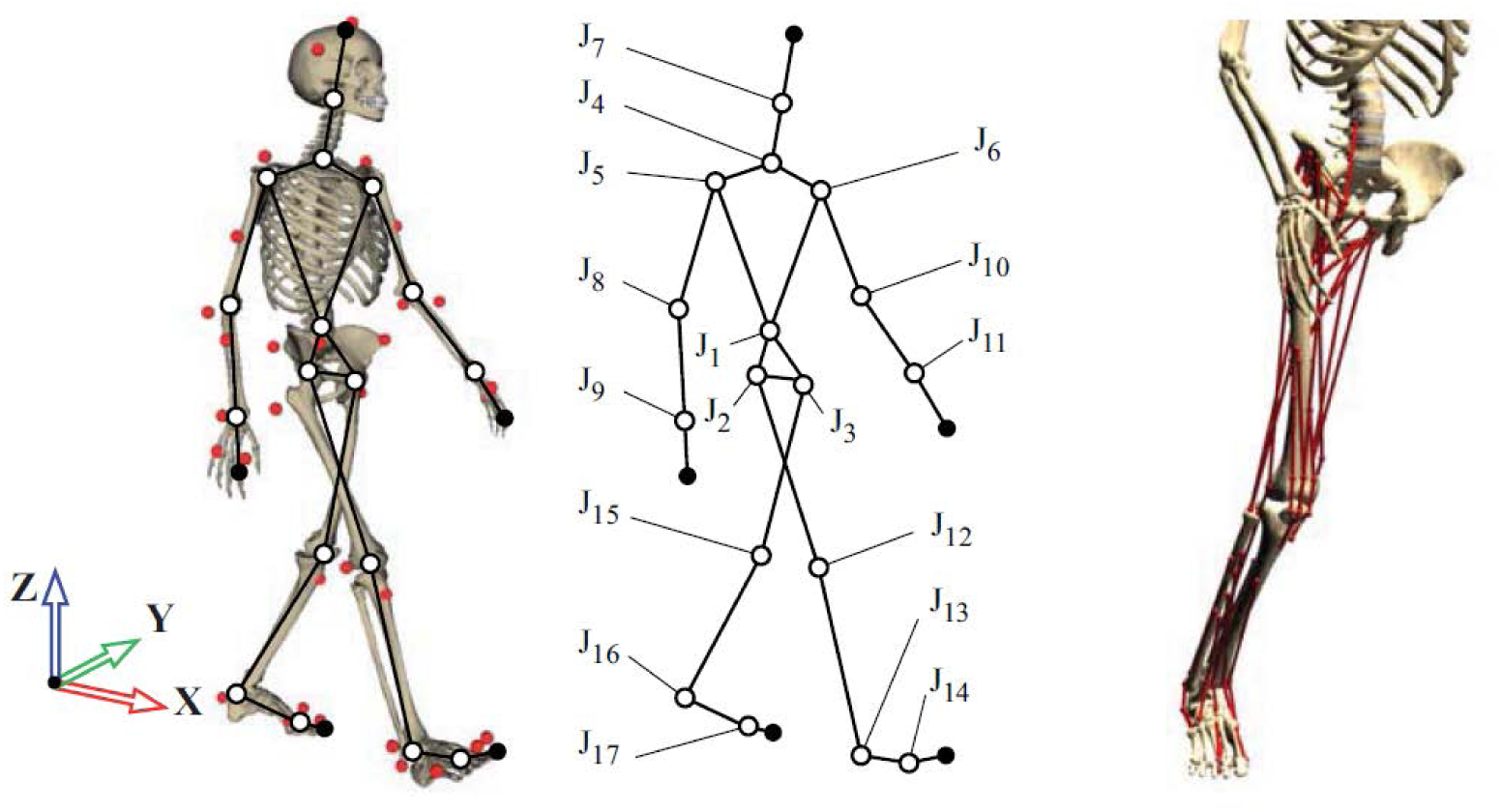
3D human model and detail of muscles on the right leg.

Matrix-R formulation (Jalon and Bayo 1994) was applied to obtain the joint torques along the motion using the in-house developed MBSLIM library (Dopico 2016) programmed in FORTRAN, as described in (Lugrís, Carlín, Luaces, et al. 2013). Once the joint torques were computed, we assumed that 43 right leg muscles contributed to six right leg inverse dynamics moments: 3 rotational DOFs at the hip, the flexion/extension DOF at the knee, and the plantar/dorsi flexion and internal/external rotation at the ankle. Muscles were modeled as one or more straight-line segments with via points. These points corresponded to the attachments of muscle and tendon to bone and were defined as the origin (i.e., proximal attachment) and insertion (i.e., distal attachment). Muscle properties and local coordinates for these points were obtained from OpenSim (model Gait2392) (Delp et al. 2007) and scaled to each subject from the generic reference OpenSim model. Length parameters (optimal muscle fiber length and tendon slack length) were scaled, for each muscle, with a scale factor calculated as the relation between the subject’s musculo-tendon length in a standing position and that of the generic model in the same position. Muscle forces were calculated from optimization-predicted muscle activations using a custom Hill-type rigid-tendon muscle model (Zajac 1989) developed in Matlab (De Groote et al. 2016). We assumed that not calibrating the positions and orientations of the joint functional axes in the leg model likely affected inverse dynamics joint moment calculations (Reinbolt et al. 2007), which in turn likely affected muscle activation calculations. Moreover, not having a process for calibrating Hill-type muscle-tendon model properties likely affected the estimated muscle activations (Serrancolí et al. 2016). However, all the methods proposed in this work were used with the same limitations.

### 2.3 Synergy Optimization

The synergy optimization (SynO) approach used in (S. Shourijeh and Fregly 2019) estimates muscle forces during human walking using synergy-constructed muscle activations, similar to the more complex approach in (Gopalakrishnan, Modenese, and Phillips 2014). In SynO, synergies couple muscle activations across time frames, requiring the optimization to be performed over all the time frames simultaneously.

Muscle synergy quantities were used as the design variables for synergy optimization. Each muscle activation synergy was composed of a single time-varying synergy activation defined by *p = (f-1)/5+1* (nearest integer, *f* = number of frames) B-spline nodal points along with its corresponding time-invariant synergy vector defined by *m* = 43 weights specifying inter-muscle activation coupling. Thus, for *n_S_* synergies (*n_S_* = 2 through 6), the number of design variables was *n_S_*(p+m)*. Each optimization problem was theoretically over-determined. However, in practice, the problems remained under-determined since neighboring time frames are not completely independent from one another.

Using these design variables, the SynO cost function was formulated as follows:

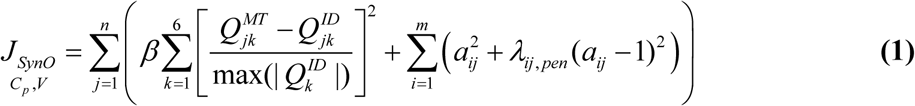

Where 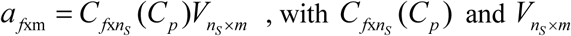 respectively the time-varying synergy activation defined by the B-spline and the corresponding time-invariant synergy vector, and 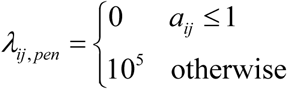. *β* = 100 is a scale factor to give more importance to the minimization of the error between 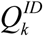 the vector of the inverse dynamics joint moments for the *k*^th^ DOF, and, 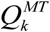 the the joint moments produced by the muscle forces estimated by SynO.

The objective function was programmed as a Fortran mex file to reduce computation time (16 times faster than the original Matlab function). Linear equality constraints made the sum of weights within each synergy vector equal to one, which made the synergy construction unique, while lower bound constraints made the synergy activation B-spline nodes and synergy vector weights non-negative. Synergy optimization problems were solved using Matlab’s *fmincon* nonlinear constrained optimization algorithm (MathWorks, Natick, MA). Five global optimizations were run using Matlab’s *ga* genetic optimization algorithm with a population size of 50, providing random initial guesses for fmincon. The SynO’s solution with the lowest objective function value was chosen as the final solution.

SynO finds muscle forces that match the inverse-dynamics joint moments as closely as possible through the moment tracking error term in the cost function. The total variance account (VAF) was used to quantify errors in inverse-dynamics joint moment matching. In contrast, the best pattern’s correlations between muscle activation and EMG were found by cross correlation with time delay using the correlation coefficient r (Matlab’s function *corrcoef*) and a maximum delay of 100 ms (S. Shourijeh et al. 2016).

### 2.4 Static Optimization

In contrast to SynO, SO’s muscle activations are independent between time frames, allowing the optimization to be performed one time frame at a time. SO was run for the same conditions as SynO (Figure 2), using the same solver *fmincon* and carrying out five global optimizations to obtain the initial guess for the initial time (then, for the remaining time-points, the initial guess is taken as the optimization result of the previous time point with the same criterion to minimize the muscle activity (sum of squares of muscle activations). Unlike SynO, SO finds muscle forces that perfectly reproduce the inverse-dynamics joint moments (in the absence of reserve actuators) through equality constraints.

**Figure 2:**
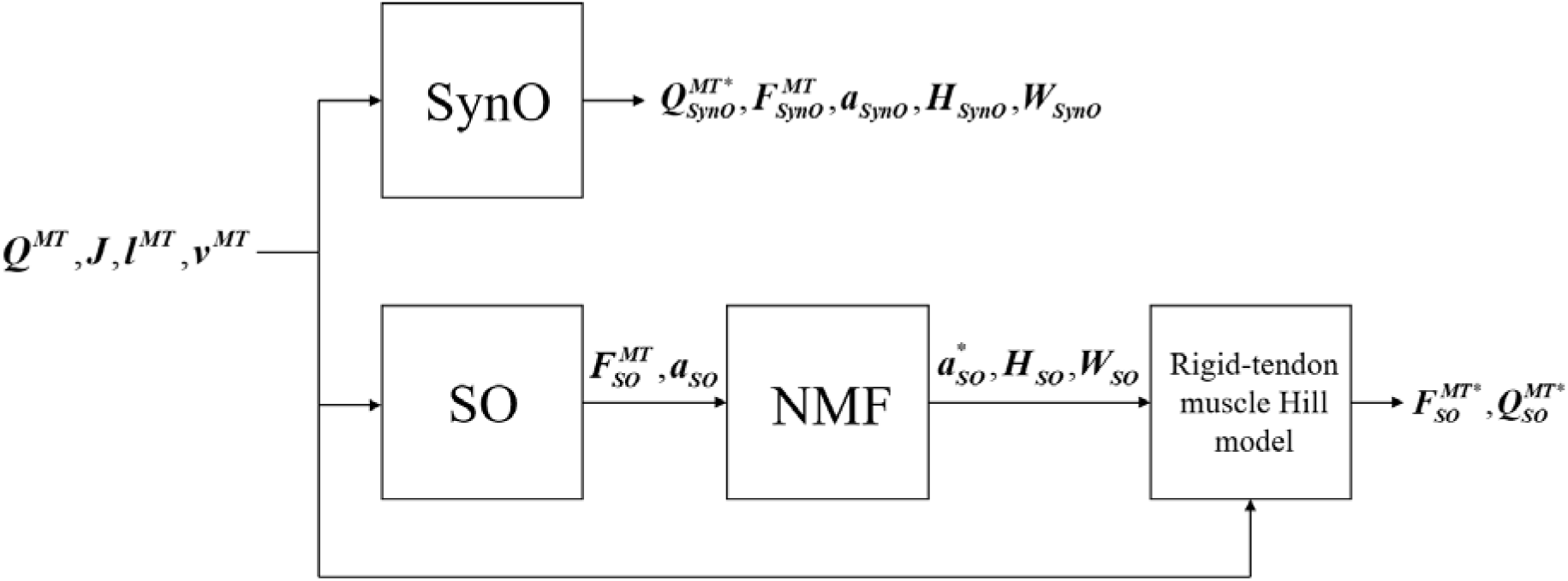
Block diagram of SynO and combined SO-NMF approaches. ***Q^MT^*** is the vector of the intersegmental moments driven by muscles, ***J*** is the Jacobian matrix of moment arms, ***l^MT^*** and ***v^MT^*** are, respectively, the length and velocity of the musculotendons. ***F^MT^*** and ***a*** represent the estimated muscular forces and activations, ***H*** the single time-varying synergy activation and ***W*** the time-invariant synergy vector. ***Q^MT*^***, ***F^MT*^*** and ***a***\* are the reconstructed intersegmental moments, muscular forces and activations.

### 2.5 Identification of muscle synergies from Static Optimization

To extract a synergy structure from the SO results, we used non-negative matrix factorization (NMF) to decompose the 43 muscles activations estimated by SO:

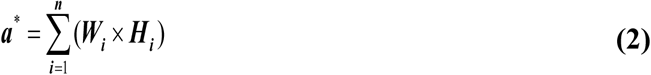

where ***a***\* is the vector of the reconstructed muscular activations, ***H_i_*** is the single time-varying synergy activation, and ***W_i_*** the corresponding time-invariant synergy vector for each of *n* synergies (*n* = 2 through 6). MATLAB *nnmf* was modified to constrain the norm-1 of each synergy vector to one to have the same constraint as SynO. Finally, using the rigid tendon Hill-type muscle model, the reconstructed muscle forces and corresponding intersegmental joint moments were derived from ***a***\*. This approach was called SO-NMF in this work.

## 3. Results

The joint moments obtained from SynO using 2 through 6 synergies matched the inverse dynamics joint moments well (Table 1 and Figure 3). The worst match was produced when using only 2 synergies, though the model was still able to match the inverse dynamics joint moments closely (mean VAF of 90%). With 3 synergies, the mean VAF obtained was higher than 96% for all the subjects. Between 4 and 6 synergies, VAF values were 98% or higher.

**Figure 3:**
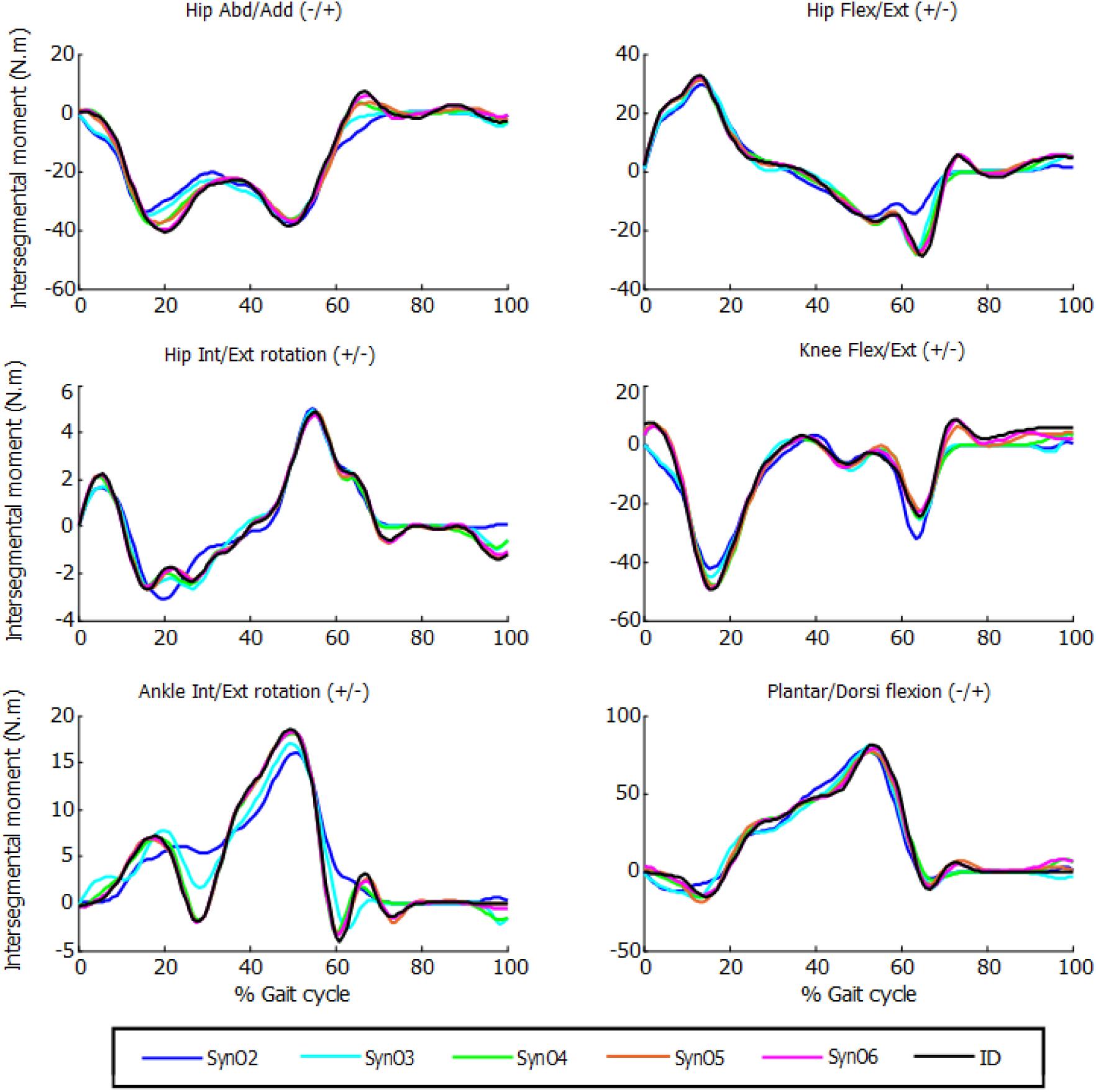
Intersegmental moments from SynO for *n* synergies (*n* = 2 through 6) vs intersegmental moments calculated by inverse-dynamics for one subject.

**Table 1:** Mean correlation VAF values across subjects between intersegmental moments calculated by inverse-dynamics and: i) joint intersegmental moments from SynO, ii) joint intersegmental moments from NMF with SO, for *n* synergies (*n* = 2 through 6) for the 5 subjects. (VAF<95% are considered not good enough, in red).

|  | Mean VAF values across subjects for joint intersegmental moment matching |  |  |  |  |  |  |  |  |  |  |
| --- | --- | --- | --- | --- | --- | --- | --- | --- | --- | --- | --- |
|  | 2 Synergies |  | 3 Synergies |  | 4 Synergies |  | 5 Synergies |  | 6 Synergies |  | SO |
|  | SynO | SO-NMF | SynO | SO-NMF | SynO | SO-NMF | SynO | SO-NMF | SynO | SO-NMF |  |
| Hip Abd/Add. | 91,25 | 73,36 | 96,63 | 86,19 | 98,91 | 85,31 | 99,39 | 92,59 | 99,79 | 92,73 | 100,00 |
| Hip Flex/Ext. | 92,99 | 78,16 | 97,90 | 94,29 | 98,95 | 95,13 | 99,70 | 93,76 | 99,80 | 97,67 | 100,00 |
| Hip Int/Ext rot. | 94,45 | 24,99 | 97,83 | 26,84 | 98,80 | 35,75 | 99,73 | 54,08 | 99,85 | 49,05 | 100,00 |
| Knee Flex/Ext. | 92,80 | 35,61 | 96,42 | 52,74 | 98,19 | 88,62 | 99,38 | 87,62 | 99,70 | 92,50 | 100,00 |
| Ankle Int/Ext rot. | 75,59 | 52,64 | 91,99 | 51,59 | 98,29 | 68,74 | 99,81 | 80,12 | 99,87 | 85,59 | 100,00 |
| Plantar/dorsi flex. | 91,87 | 75,04 | 95,80 | 80,70 | 98,17 | 84,98 | 98,93 | 91,69 | 99,69 | 91,02 | 100,00 |
| **Mean Across Joints** | **89,82** | **56,63** | **96,10** | **65,39** | **98,55** | **76,42** | **99,49** | **83,31** | **99,78** | **84,76** | **100,00** |

While SO exactly reproduced the inverse dynamics joint moments through its equality constraints, SO-NMF’s muscular activations with 2 through 6 synergies matched the experimental inverse dynamic joint moments poorly (Table 1 and Figure 4). With 2 and 3 synergies, matches for some joint moments were worse than 50% VAF, and the mean match was lower than 70%. Between 4 and 6 synergies, mean VAF values were between 76% (with 4 synergies) and 90% (with 6 synergies), and some joint moments remained under 80%.

**Figure 4:**
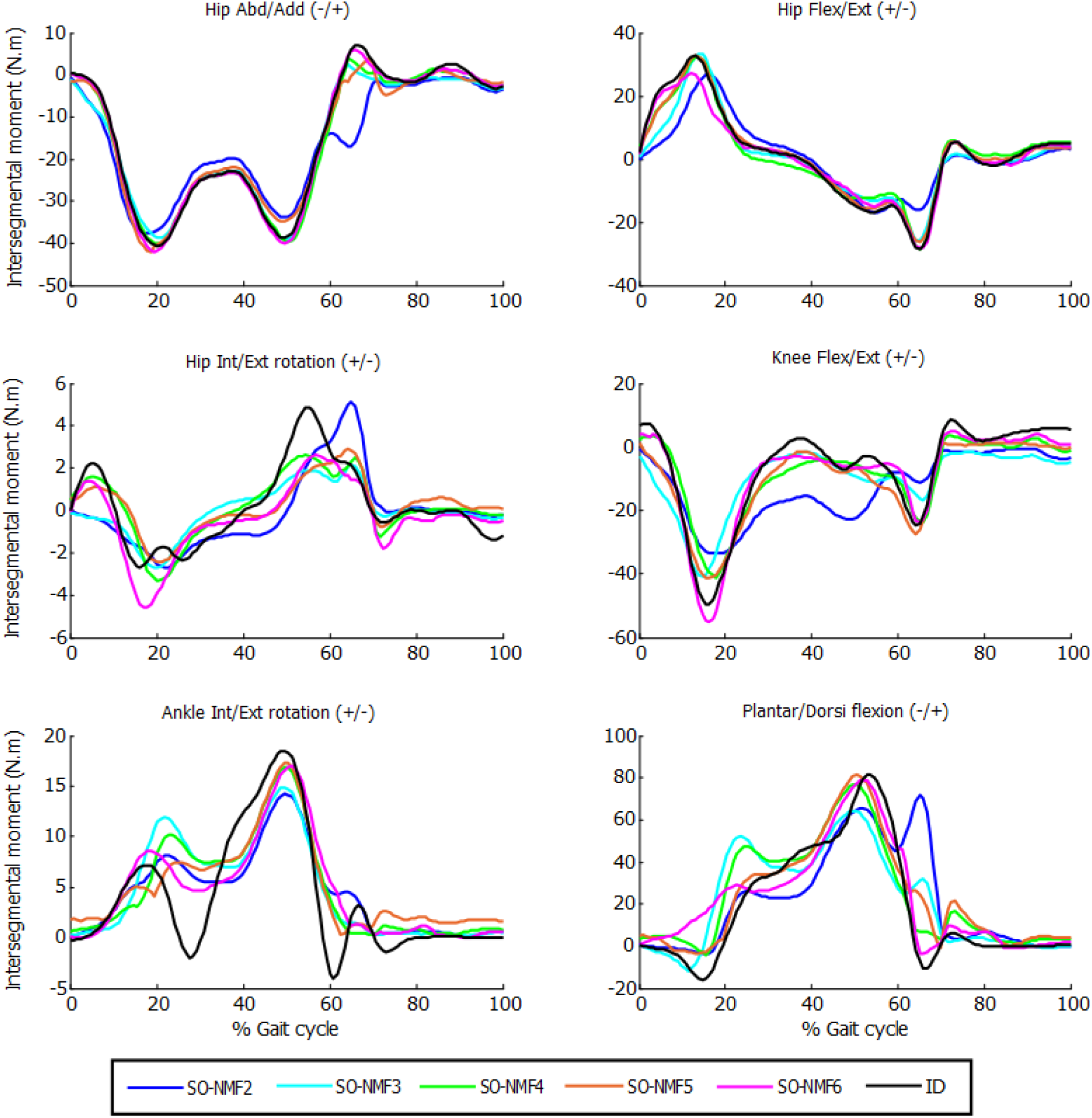
Intersegmental moments from SO and NMF for *n* synergies (*n* = 2 through 6) vs intersegmental moments calculated by inverse-dynamics for one subject.

Comparison of muscle activations estimated using SynO with experimental EMG measurements showed significant differences when the number of synergies was increased (example in Figure 5 for one of the subjects). Activations estimated by SynO became more similar to those estimated by SO as the number of synergies was increased. However, the mean correlations r between estimated muscle activations and measured EMG patterns for the five subjects did not present such differences (Table 2). Mean values of the different approaches were close, between 0.56 (4 synergies) and 0.62 (6 synergies) for SynO and 0.60 for SO.

**Figure 5:**
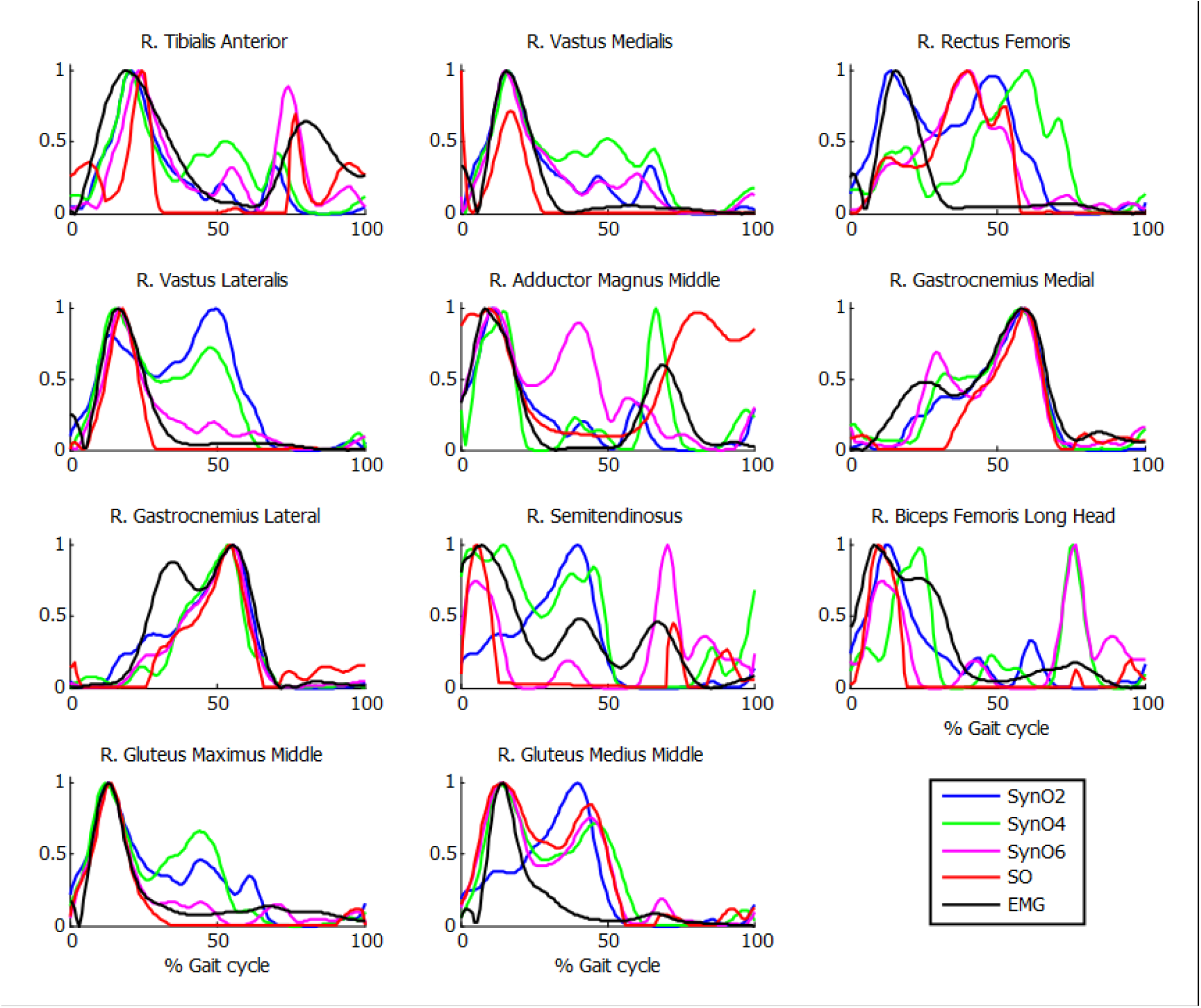
Normalized muscle activations obtained for one subject from SynO and *n* synergies (*n* = 2 through 6) vs normalized EMG.

**Table 2:** Mean across subjects correlation coefficient r values between EMG measurements and: i) muscular activations from SynO, ii) muscular activations from NMF with SO, for *n* synergies (*n* = 2 through 6) of the 5 subjects (r<0.4, in red, are considered poor and r≥0.6, in green, are considered good).

|  | Pearson corr. Coeff. r between across-subject mean EMG vs. Muscle activations |  |  |  |  |  |  |  |  |  |  |
| --- | --- | --- | --- | --- | --- | --- | --- | --- | --- | --- | --- |
|  | 2 Synergies |  | 3 Synergies |  | 4 Synergies |  | 5 Synergies |  | 6 Synergies |  | SO |
|  | SynO | SO-NMF | SynO | SO-NMF | SynO | SO-NMF | SynO | SO-NMF | SynO | SO-NMF |  |
| R. Tibialis Anterior | 0,58 | 0,52 | 0,59 | 0,75 | 0,56 | 0,75 | 0,56 | 0,74 | 0,69 | 0,76 | 0,69 |
| R. Vastus Medialis | 0,84 | 0,74 | 0,61 | 0,84 | 0,68 | 0,74 | 0,68 | 0,75 | 0,73 | 0,76 | 0,71 |
| R. Rectus Femoris | 0,53 | -0,07 | 0,52 | -0,09 | 0,25 | -0,02 | 0,08 | -0,06 | 0,16 | -0,05 | -0,04 |
| R. Vastus Lateralis | 0,72 | 0,74 | 0,75 | 0,82 | 0,65 | 0,80 | 0,54 | 0,78 | 0,73 | 0,79 | 0,74 |
| R. Adductor Magnus Middle | 0,48 | 0,54 | 0,43 | 0,71 | 0,52 | 0,51 | 0,57 | 0,56 | 0,57 | 0,56 | 0,52 |
| R. Gastrocnemius Medial | 0,72 | 0,80 | 0,87 | 0,71 | 0,77 | 0,67 | 0,70 | 0,69 | 0,75 | 0,67 | 0,60 |
| R. Gastrocnemius Lateral | 0,57 | 0,72 | 0,76 | 0,77 | 0,67 | 0,66 | 0,70 | 0,70 | 0,64 | 0,71 | 0,57 |
| R. Semitendinosus | 0,36 | 0,73 | 0,58 | 0,89 | 0,53 | 0,66 | 0,66 | 0,67 | 0,50 | 0,60 | 0,57 |
| R. Biceps Femoris Long Head | 0,72 | 0,69 | 0,57 | 0,85 | 0,51 | 0,79 | 0,50 | 0,80 | 0,54 | 0,86 | 0,84 |
| R. Gluteus Maximus Middle | 0,74 | 0,71 | 0,71 | 0,86 | 0,71 | 0,90 | 0,84 | 0,92 | 0,87 | 0,92 | 0,91 |
| R. Gluteus Medius Middle | 0,25 | 0,39 | 0,45 | 0,41 | 0,36 | 0,40 | 0,48 | 0,42 | 0,38 | 0,43 | 0,44 |
| **Mean** | **0,59** | **0,60** | **0,62** | **0,68** | **0,56** | **0,62** | **0,57** | **0,64** | **0,59** | **0,64** | **0,60** |

Reconstructed muscle activations obtained using SO-NMF poorly matched the activations estimated using SO (Table 3 and Figure 6). Using only 2 synergies, a mean r^2^ correlation of 0.44 was obtained for the 43 muscles, and a maximum correlation of 0.87 was obtained with 6 synergies. However, while reconstructed muscle activations and reconstructed joint moments showed low correlations with SO results, correlations between experimental EMG patterns and the newly reconstructed activations showed better mean values. The best correlations were obtained using 3 synergies, with a mean value of 68%. From 2 to 6 synergies, the correlations varied between 60 and 68%, giving similar or better results than those obtained using SO estimated activations.

**Figure 6:**
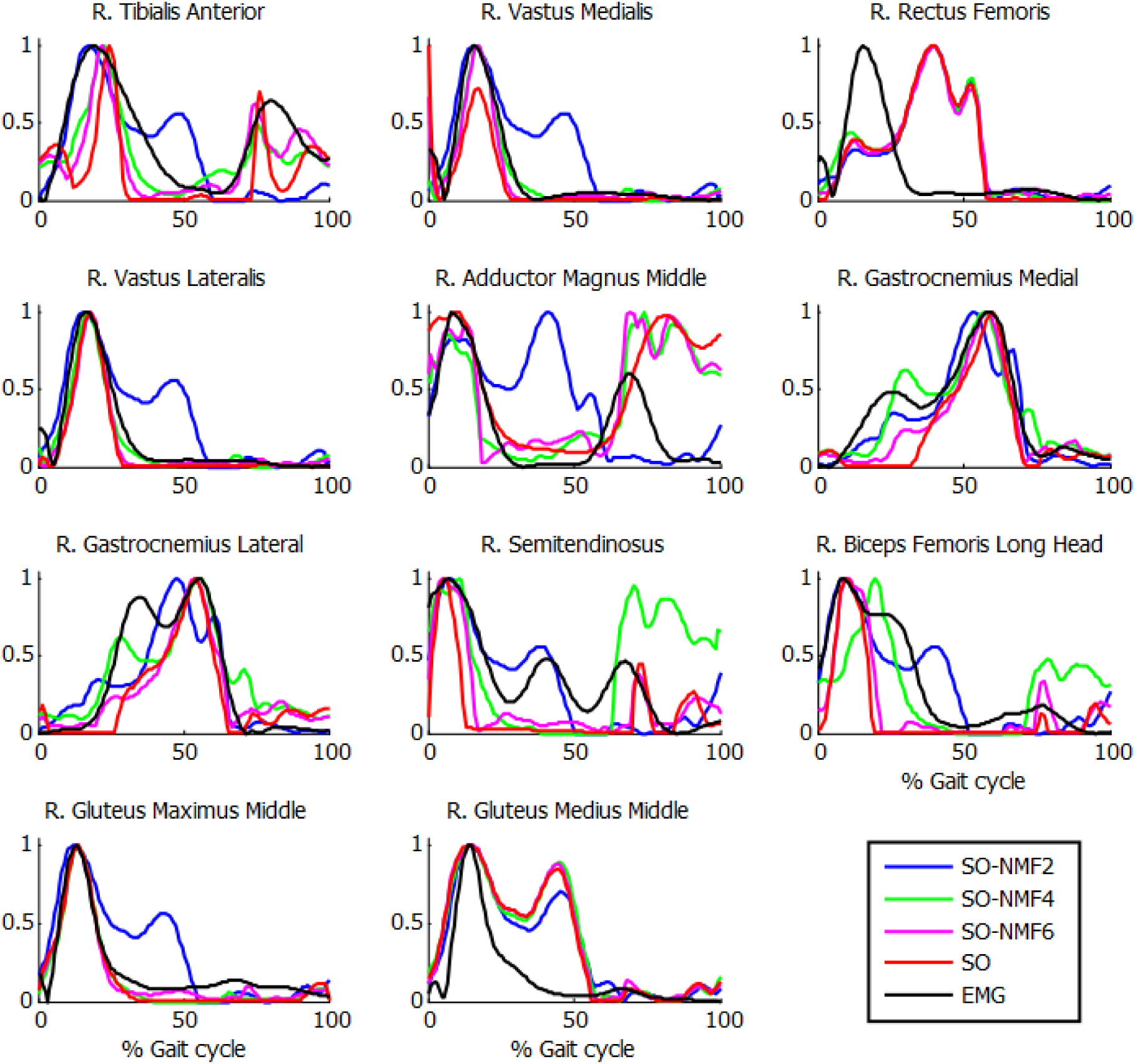
Normalized muscle activations obtained for one subject from SO and NMF with *n* synergies (*n* = 2 through 6) vs normalized EMG.

**Table 3:** Mean correlation coefficient R^2^ values between muscular activations calculated by SO and SO-NMF for *n* synergies (*n* = 2 through 6).

|  | **r<sup>2</sup> mean values between SO and SO-NMF** |  |  |  |  |
| --- | --- | --- | --- | --- | --- |
|  | **2 Synergies** | **3 Synergies** | **4 Synergies** | **5 Synergies** | **6 Synergies** |
| ***a* vs. *a*<sup>\*</sup>** | 0,44 | 0,56 | 0,75 | 0,83 | 0,87 |

Finally, the computational efficiency of the different approaches studied in this work was compared in Table 4. All calculations were performed on an Intel® Core™ i7-6700K processing running at 4.00 GHz with 16 GB of RAM, and all functions (except the objective function of SynO programmed in a *mex* file) were programmed in Matlab using the optimization function *fmincon* without parallelization. Computation time increased significantly with the number of synergies, and with SO clearly being the fastest method, requiring a mean duration of 2 s to solve a complete gait cycle of approximately 1 s. The NMF analysis required about 1 s in Matlab.

**Table 4:** Mean computational time for SO and SynO with *n* synergies (*n* = 2 through 6) of the 5 subjects.

|  | SynO2 | SynO3 | SynO4 | SynO5 | SynO6 | SO |
| --- | --- | --- | --- | --- | --- | --- |
| Computational time (sec) | 80 | 115 | 196 | 317 | 587 | 2 |

## 4. Discussion

This work analyzed if a recent synergy-based approach used to solve the muscle force sharing problem, called SynO (S. Shourijeh and Fregly 2019), can improve estimation of muscle activations during gait. In addition to comparing the correlations between estimated activations obtained by SO and SynO for five healthy subjects, we explored the reliability of predicting muscle activations by applying NMF to SO’s muscle activations. Increasing the number of synergies from 2 until 6 in SynO had minimal influence on the model’s ability to match inverse dynamics joint moments closely. On the other hand, reconstructed joint moments from SO combined with NMF matched inverse dynamic joint moments poorly, since unlike SynO, NMF does not take into account any joint moment information. Consequently, the resulting joint moments would produce a new motion, different from the original one.

Muscle activations obtained from SynO using 2 through 6 synergies presented visual differences in patterns, which explain why the joint stiffness’s estimated in (S. Shourijeh and Fregly 2019) were affected by the number of synergies. Increasing the number of synergies implies increasing the number of design variables, thus allowing more freedom in the behavior of muscle activations. For this reason, SO presented results closer to SynO with six synergies. The same observations can be made with NMF when varying the number of synergies (Figure 6).

The highest muscle activations were observed for two synergies (blue line), which generated higher co-contraction when seeking to match the intersegmental moments, which would likely produce higher joint stiffness (S. Shourijeh and Fregly 2019). Individuals with neurological disorders such as stroke or Parkinson’s disease often use a lower number of muscle synergies than do healthy individuals (Clark et al. 2009; Rodriguez et al. 2013). Consequently, individuals with these disorders may generate higher stiffness to maintain stability and reject walking disturbances (Kitatani et al. 2016; Rinalduzzi et al. 2015).

Correlations observed in Table 2 are reasonable in general, with mean *r* values for the five subjects varying between 56% and 68%. Surprisingly, no significant differences were observed for different numbers of synergies. The poorest results were obtained for the rectus femoris and the gluteus medius. Crosstalk (Jungtäubl et al. 2018) may explain the low correlation for these muscles, especially rectus femoris. Comparing the rectus femoris EMG signal with the vastus intermedius (muscle located under the rectus femoris) estimated activation resulted in a higher correlation (from 0,25 to 0,61).

Strangely, the reconstructed activations from SO-NMF matched EMG better than did the original activations from SO. However, the reconstructed inverse dynamics joint moments showed a poor correlation VAF (between 56% and 85%), thus producing an inconsistent actuation. This might have been caused by the use of a reduced number of components when obtaining the synergy information through NMF for a large number of muscles.

For SynO as well as for SO-NMF, the best correlations with experimental EMG patterns were obtained using three synergies. As mean intersegmental moment matching with three synergies was good using SynO (96.1% in Table 1, although the matching of the internal/external rotation moment at ankle was only of 92.0%), it appears that the central nervous system (CNS) could control one leg during gait using only three synergies. Olree (Olree and Vaughan 1995) recorded EMG signals bilaterally from eight leg muscles and also showed that three basic patterns could account for the locomotion activity of these muscles. However, based on EMG activity analysis of 16 unilateral leg muscles (Winter and Yack 1987), Davis (Davis and Vaughan 1993) and Ivanenko (Ivanenko, Poppele, and Lacquaniti 2004) concluded, respectively, that four and five patterns could be necessary. As explained in (Banks et al. 2017) and (Steele, Tresch, and Perreault 2013), variations in methodological choices, as unilateral or bilateral analysis, selected muscles, EMG processing, or computational method may generate different results. This, it is difficult to conclude what number of synergies is used by the CNS during gait. In this work, though only one leg was studied, it would be interesting to explore how many bilateral synergies would be found using SynO when studying both legs together, especially in the case of unilateral stroke (Sainburg, Good, and Przybyla 2013; Coscia et al. 2015).

Comparing EMG correlations obtained with SynO and *n* synergies (*n* = 2 through 6) and those obtained with SO, there are no clear advantages between the two approaches. Despite its higher dimensional control space, SO presented a mean r correlation of 0.60 with the experimental data. SynO produced better correlations only for the 3 synergy case.

In conclusion, this study evaluated the ability of the SynO approach to predict muscle activations obtained from experimental EMG measurements during gait and found that three synergies are theoretically enough to control leg muscles during gait. However, no significant differences in ability to predict experimental EMG patterns were found between SynO with *n* synergies (*n* = 2 through 6) and SO, so thus neither approach can be considered preferable for this purpose. While SO is computationally faster and requires muscle forces to match inverse dynamic joint moments through constraints, extraction of synergies by NMF from SO’s results generated new intersegmental joint moments that were inconsistent with the experimental joint moments. The SynO approach offers reasonable prediction of muscle activations using an imposed synergy structure and reduced dimensional control space and could be useful for applications such as functional electrical stimulation and motion control and prediction.

## Acknowledgments

This work was funded by the Spanish MINECO under project DPI2015-65959-C3-1-R, co-financed by the EU through the EFRD program, and by the Galician Government under grant ED431B2016/031. Moreover, F. Michaud would like to acknowledge the support of the Spanish MINECO by means of the doctoral research contract BES-2016-076901, co-financed by the EU through the ESF program.

